# Tau pathology leads to lonely non-traveling slow waves that mediate human memory impairment

**DOI:** 10.1101/2024.05.22.595043

**Authors:** Omer Sharon, Xi Chen, Jason Dude, Vyoma D. Shah, Yo-El S. Ju, Willam J. Jagust, Matthew P Walker

## Abstract

Memory markedly declines with age, exaggerated in those with Alzheimer’s disease, yet the mechanisms are still not resolved. Here, we show that frontal lobe tau pathology in humans leads to impaired en masse unity and cortical traveling propagation of NREM slow waves, consequentially impairing memory retention. We elucidate these findings using PET tau brain imaging, and then replicate and extend them using AD pathology markers derived from lumbar puncture CSF in an independent clinical cohort. Thus, tau-associated memory deficits are not wholly direct, but indirectly mediated through consequential “lonely”, non-traveling slow-wave events.

## Main

Brain aging is characterized by a decline in memory function, including the ability to retain new long-term episodic memories^1^, though the mechanistic reasons underlying such impairment are not yet fully understood. Moreover, such memory impairment is exaggerated in those with Alzheimer’s disease (AD) and the associated pathology^2,3^. Specifically, human and rodent findings continue to implicate aggregates of tau protein (as neurofibrillary tangles) in particular as a pathological feature consistently predicting memory dysfunction cross-sectionally, and longitudinally cognitive decline^4–6^. Indeed, the degree of age-related memory impairment varies across individuals^7,8^, suggesting that memory decline is age-*related* but not age-*dependent*. This has led to the proposal that other unrecognized factors contribute to cognitive decline associated with aging and AD pathology^9^.

One potential explanatory candidate is sleep impairment. To date, amyloid-beta (Aβ) and tau proteins have been associated with deficits in classic features of sleep, notably a diminution in non-REM (NREM) slow-wave sleep, and the number and size (amplitude) of NREM slow waves linked to impaired memory^10^. However, these classical sleep oscillation features only explain a portion of the variance in cognitive impairment, leaving unknown other contributing factors.

Of relevance, electrophysiological studies have discovered that NREM slow waves are not solely characterized by their number or amplitude, nor integrated power. Instead, slow-wave oscillations are dynamic, and dynamic in at least two ways. First, NREM slow waves appear in a coordinated en masse ensemble across multiple sites, demonstrating that slow waves form a collective unity across large cortical territory^11–14^. Second, slow waves are not stationary, but instead, traverse across the cortex in long-range traveling paths, most typically from prefrontal origins^11,14–16^.

While these physiological characteristics have recently been described, to date no functional benefit(s) has been ascribed to either of these contemporary features of NREM slow oscillations in healthy individuals or disease states. Moreover, how regional tau burden in aging impacts the two novel features of NREM slow waves remains uncharacterized, as does any relationship with failed memory consolidation leading to overnight forgetting rather than remembering.

Building on these findings, and using two different independent cohorts and two different measures of tau pathology burden, the current studies sought to test three interrelated hypotheses: 1) brain aging impairs two dynamic properties of slow wave oscillations: (i) their ensemble coordinated activity, and (ii) their traveling wave propagation, 2) the magnitude of these impairments in older adults is not age-dependent, but pathology dependent, specifically predicted by the severity of tau burden in slow-wave-generating regions of the frontal lobe, and finally 3) in older adults, tau-associated memory deficits in the aging human brain are not simply direct, but also partially mediated through impairments in the ensemble coordinated activity and traveling wave propagation.

In brief (and see Methods), two independent cohorts of adults went through multi-night and day experiments involving in-lab overnight polysomnographic multichannel EEG scalp. Data from the first cohort was collected at UC Berkeley (N=135) and composed of both young (N=61, mean age=20.24, SD=1.91, 49% male) and older adults (N=74, mean age= 74.79, SD= 5.73, 32% male) from the Berkeley Aging Cohort Study (BACS). The older adults had a range of tau protein and beta-amyloid measured with PET imaging of [18F]Flortaucipir (FTP) and [11C]PIB, respectively, and who were not in the stages of mild cognitive impairment (MCI) nor dementia. In addition, in the BACS cohort, a sleep-dependent episodic memory task was performed before and after the overnight EEG recordings in all older adults and some young adults (N=16/61). Data from the second cohort was collected through the “Biomarkers of Alzheimer disease in Sleep and Electroencephalography” (BASE) study at Washington University in St Louis, and was exclusively comprised of older individuals (N=81 mean age = 70.5 SD=4.4 years, 48% males), with paired CSF amyloid and tau measures, and with psychometric memory performance assessment (see methods for details).

First, analyses focused on testing whether older adults suffered impaired en masse cortical wave coordination, and impaired cortical traveling wave propagation, relative to young adults (comparing both age groups in the Berkeley cohort). Individual slow wave oscillation events were detected (slow oscillations, 0.3–1.5 Hz) using established algorithms^11,17,18^. Thereafter, analyses focused on the two a priori experimental metrics: 1) the degree of cortical involvement of the slow oscillations, identifying whether the slow oscillation appeared in an ensemble, collective manner, or instead, was spatially isolated, reflecting a more “lonely” nature of slow waves, and 2) the cortical traveling nature of the slow waves, indexed using the distance traveled by every slow oscillation (see **Fig 1** and Methods).

**Fig 1.**
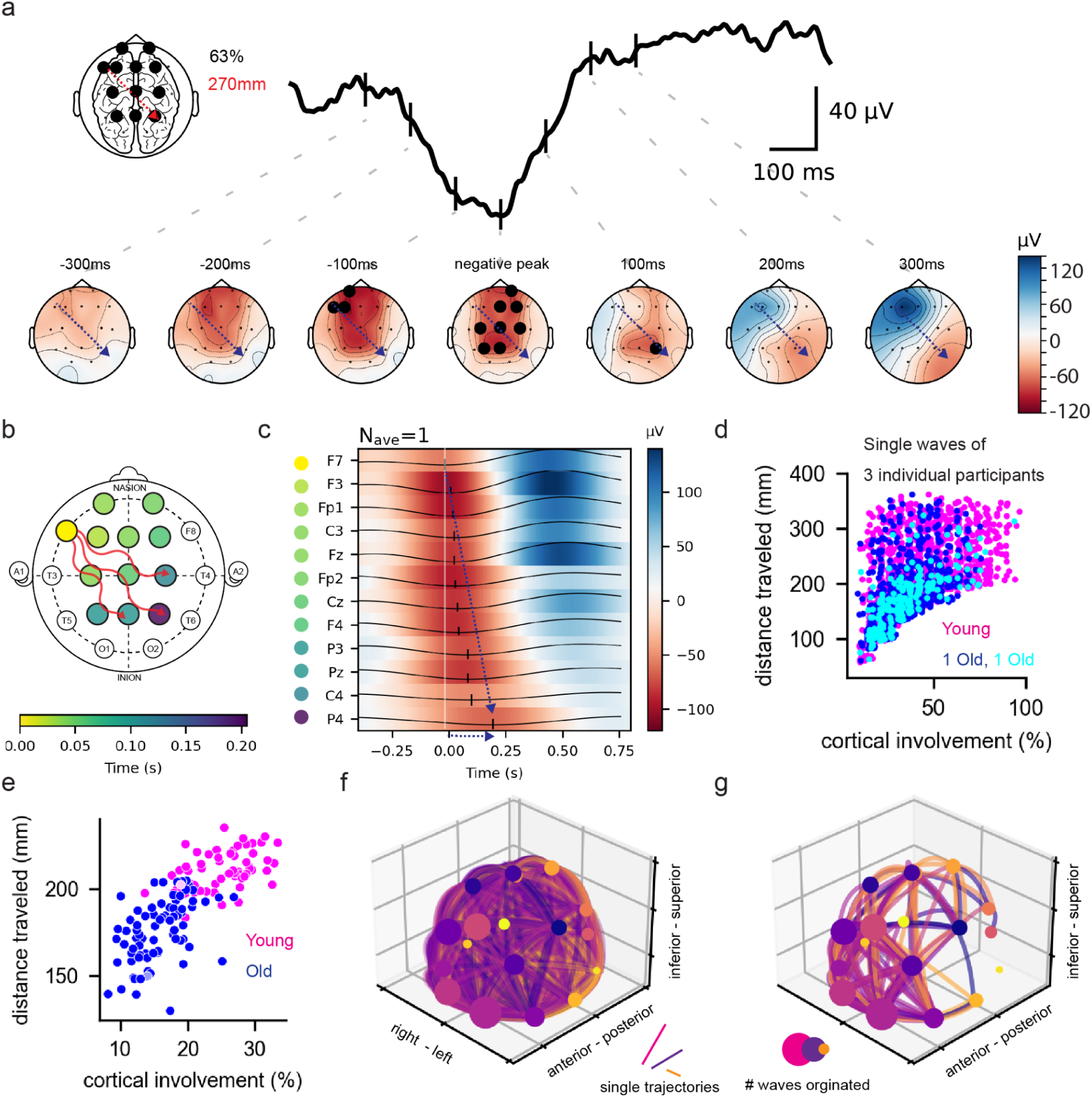
Traveling Slow Oscillations detected in young and older adults’ clinical EEG. (a) One single traveling slow oscillation averaged across all electrodes (black trace) and the voltage across the scalp (below), colored in black are electrodes that crossed the detection threshold, the slow wave direction from the origin to the furthest electrode is depicted by the black arrow. The inset shows the resulting two metrics further used across participants and groups demonstrated in relation to the example: cortical involvement (the number of black electrodes in the inset divided by the total number of electrodes = 63%), and the distance traveled (red arrow = 270mm, inset) that is the length of the trajectory from the electrode of origin (yellow in b) to the furthest electrode (dark blue in **b**). (**b**) Illustration of the electrode location and time-flow of one single traveling slow oscillation (same as in a) across the cortex. (c) Time domain amplitude heatmap of colored electrodes in **a**, showing the advancement of the negative peak in time. (dashed arrow). (**d**) Cortical involvement and distance traveled of single waves within 3 individual subjects, one young (magenta) and two older adults (dark and light blue). Note that within every level of cortical involvement different single slow oscillations can show different degrees of travel, 1% of random noise was added in both dimensions to avoid overlap (**e**) Average data across single subjects, young (magenta) and older (blue). (f) Traveling slow waves in a young individual participant marked with a lighter color in e. The size of each dot represents the number of waves originating from the respective electrode, while the line colored by the same color of its origin represents the detected outgoing trajectory of a single slow oscillation across the electrodes. (**g**) Same as in **f** for a single older participant (marked in light magenta in e). Note that beyond overall changes in amplitude and quantity of slow oscillations, the older individual shows a similar number of slow oscillations in frontal electrodes but these do not travel as much as in the young individual.

Focusing on the first (of the two) slow-wave metrics of en masse ensemble cortical involvement, on average, slow oscillations were co-detected in 24.98% ±4.56% of the EEG electrodes in young healthy adults, demonstrating a robust communal, cortically collective slow-wave event process. In contrast, and consistent with the hypothesis, there was a significant impairment in the coordinated unity of slow wave in older adults showing an average cortical involvement of 15.79% ±3.91% (T=-12.35 p=<10^-23^, cohen-d=2.16) of the electrodes (**Fig 2a**). In prevalence terms, in the young adults, 27.21% ±8.30% of slow oscillations were simultaneously detected in 6 or more electrodes across the scalp, demonstrating that the ensemble slow oscillations are frequent in the young. In contrast, only 9.61% ±6.74% of slow oscillations were simultaneously detected in 6 or more electrodes in the older adults, a 3-fold impairment in older relative to young adults (U=251 p<10^-18^). Older participants also showed lower slow wave amplitude (106.28µV ±9.62µV) compared with young (134.76µV ±10.98µV, T=-15.85, p<10^-30^) as well as lower slow wave density (1.62±1.71 waves per minute) compared with young (6.17±2.35 per minute). However, these differences in cortical involvement went beyond the previously reported lower amplitude and density of slow waves in the elderly. Analysis of covariance showed that only 39.5% (F =88. p<10^-14^) of the between-participant variance in cortical involvement can be explained by amplitude and only 20% by density (F=33 p=10^-8^), indicating that differences in cortical involvement go above and beyond previously described reduced slow wave amplitude and density in elderly^19^.

**Fig 2.**
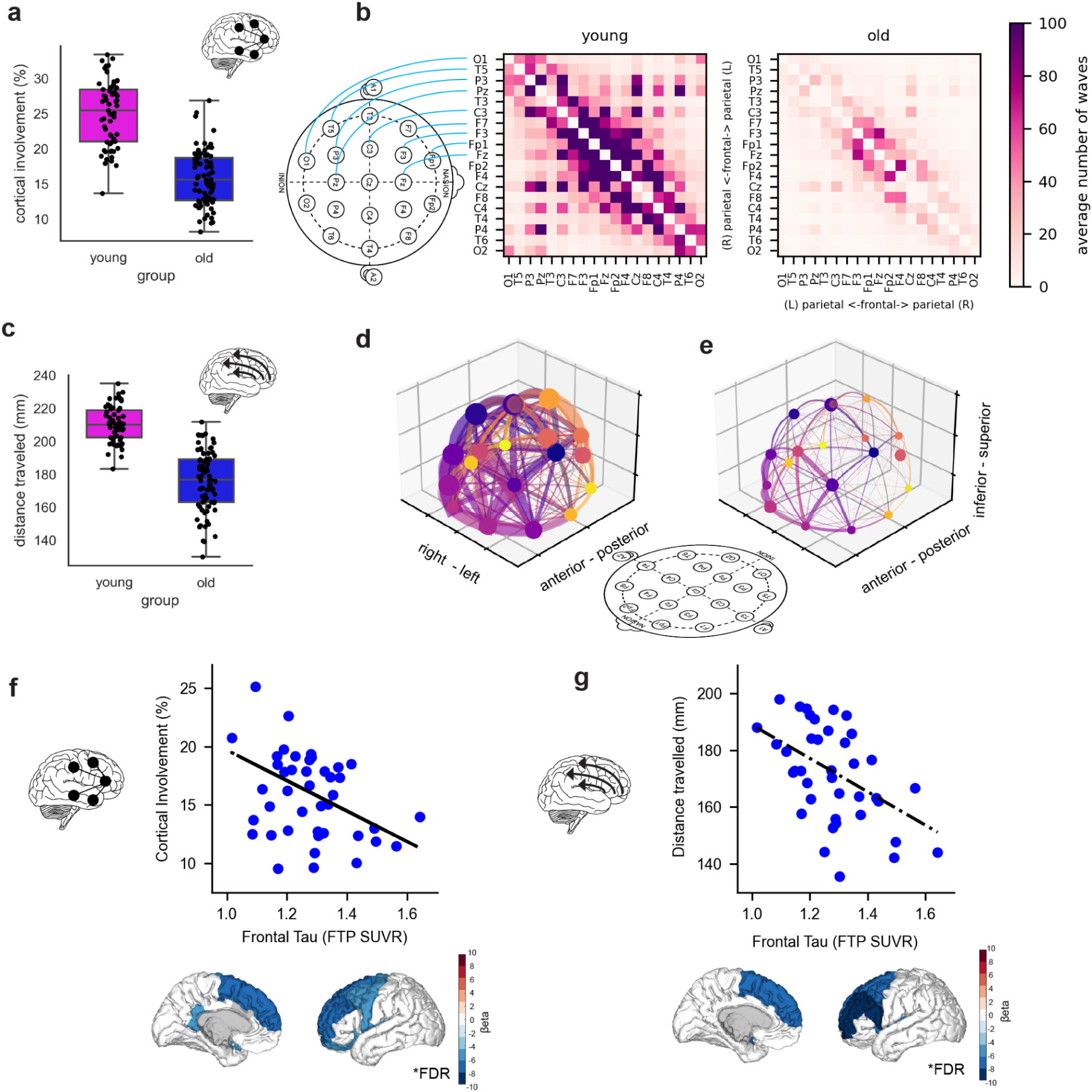
Older Adults show impaired slow oscillation travel, restricted to the frontal lobe and predicted by frontal tau accumulation. (a) Cortical involvement in slow oscillations in young (magenta) and older participants (blue) (b) The average number of waves traveling between each pair of the electrodes, the center of the matrix represents frontal electrodes while the upper part represents the left hemisphere from the front to back and the lower part the right hemisphere in the same manner (see inset on the left). Note how in the older group slow oscillations do not travel outside of the frontal areas of the brain while young participants are more equally distributed (see Fig S3 for relative terms) **(c)** Cortical distance traveled by slow oscillations in young (magenta) and older participants (blue) **(d)** The average slow wave count as represented in Fig 3b left, but in 3-dimensions. Here, the size of each dot represents the average number of waves originating from the respective electrode, while the line colored by the same color of its origin represents the average number of detected outgoing trajectories across the electrodes. **(e)** Same as in d for older participants. **(f)** Frontal tau burden in frontal ROI (see S5) predicts the degree of cortical involvement in slow oscillations, below are beta values all 34 cortical ROIs in the Desikan-Killany atlas showing specificity for frontal regions, FDR corrected **(g)** Frontal tau burden in frontal ROI (see S5) predicts the degree of distance traveled by slow oscillations. Below are beta values of all 34 cortical ROIs in the Desikan-Killany atlas showing specificity for frontal regions, FDR corrected.

Beyond aging impairing the amount of cortex involved in the mass coordinated occurrence of slow waves, the next analyses tested the hypothesis that aging further impairs the dynamic traveling, propagating nature of slow waves across the scalp territory. In young adults, not only were larger cortical territories involved but there was robust cortical traveling of slow oscillations. The average traveling scalp propagation in young was 210.1mm±10.9. In contrast, the slow waves of older adults exhibited diminished cortical travel, with a propagation traveling distance of 176.1mm±18.0. Compared with young adults, this represents a relative propagation impairment of 16.18% in older adults (T=-13.61 p<10^-25, cohen-d=2.21). Similarly to cortical involvement, analysis of covariance showed that only 28% (F=52, p<10^-10)^) of the between-participant variance in distance traveled was explained by amplitude and only 42% by slow wave density (F=42.9, p<10^-8^), establishing that differences in slow-wave travel extend beyond previously reported decreases in the amplitude and density of slow waves in the old^19^.

For both slow-wave metrics, greater chronological age in older adults was correlated with the degree of impairment in the coordinated en masse slow-wave occurrence (r^2^=-0.29, p=0.02), and the decreased traveling distances across the scalp (r^2^=-0.36 p=0.003). Thus, a degree of impairment in these two contemporary slow-wave features, when considered independent of brain pathology, would appear to be age-related. However, to address whether Alzheimer’s disease pathology provided a superior accounting for these age-related associations, analyses next examined whether *a priori* tau burden in the frontal lobe more accurately explained these impairments across older adults when including age as a covariate.

In examining slow-wave generating frontal regions of interest (see Methods), the extent of tau burden within the frontal cortex was positively and significantly related to the degree of impairment in the en masse coordinated ensemble activity. Here, and when controlling for chronological age alone we found that frontal tau predicted the decrease in cortical involvement (β=-8.91 T=-2.14 p=0.03, r^2^=0.15) but age did not (β=-0.10 T=-1.10 p=0.27, r2=0.15). However, beyond age, other factors might be confounding the effects by influencing both tau levels and slow waves. In particular, sex^20,21^, Apnea-Hypopnea Index (AHI)^22^, and the degree of amyloid-beta burden (indexed using [^11^C]PIB^10^) as well as the degree of gray matter atrophy^23^ within the frontal ROI. When including all these factors in the regression model (see Methods), frontal tau was the sole factor predicting impaired coordinated slow-wave events, leading to more isolated, “lonely” slow waves in a significant manner (β=-13.06 T=-2.59 p=0.01, r2=0.32, and see **Table 1** in extended data). Beyond the a priori frontal areas, post-hoc analyses including all areas in the Deiskan Killany atlas confirmed frontal tau is key and further revealed that tau burden in the parahippocampus and the isthmus cingulate cortex of the medial temporal lobe (MTL) also predicted impaired cortical involvement (both p<0.001, FDR corrected), of interest considering both regions have been implicated in episodic memory impairment in older adults and those with Alzheimer’s disease (See **Fig 2h** bottom) (**Fig 2g**, cortical involvement).

**Table 1.** Linear regression of frontal tau and cortical involvement.

| | $\beta$ | SE | T | p-value | CI[2.5%] | CI[97.5%] | Relative importance (%) |
| --- | --- | --- | --- | --- | --- | --- | --- |
| Intercept | 43.80 | 15.05 | 2.91 | 0.01 | 12.98 | 74.62 |  |
| frontal tau | -13.06 | 5.04 | -2.59 | 0.02 | -23.38 | -2.73 | 53.88 |
| Age | -0.20 | 0.11 | -1.92 | 0.07 | -0.42 | 0.01 | 33.82 |
| PiB | 2.01 | 3.51 | 0.57 | 0.57 | -5.18 | 9.19 | 4.51 |
| Sex (Male) | -1.05 | 1.20 | -0.87 | 0.39 | -3.51 | 1.41 | 4.33 |
| AHI | 0.04 | 0.06 | 0.66 | 0.52 | -0.08 | 0.16 | 2.51 |
| frontal thickness | 0.12 | 0.77 | 0.16 | 0.87 | -1.45 | 1.69 | 0.95 |

Travel measures were correlated with amplitude and the number of detected slow oscillations (r^2^>0.44, p<0.006 for all correlations), but are not simply derivative of each other. Artificially stratifying the data (see Methods) such that all older participants have the same median amplitude (120µV) did not prevent a significant predictive power for cortical involvement (**Fig 3c**, β=-8.37 T=-2.75 p=0.03 r2=0.14). Selecting only the first one hundred waves per subject to control for differences in the number of waves yielded similar results (**Fig 3d**, β=-11.72 T=-2.69 p=0.01 r2=0.28), suggesting deficits in the cortical involvement in slow waves and their travel contribute in a manner distinct and different to recognized decreases in slow wave density and amplitude.

**Fig 3:**
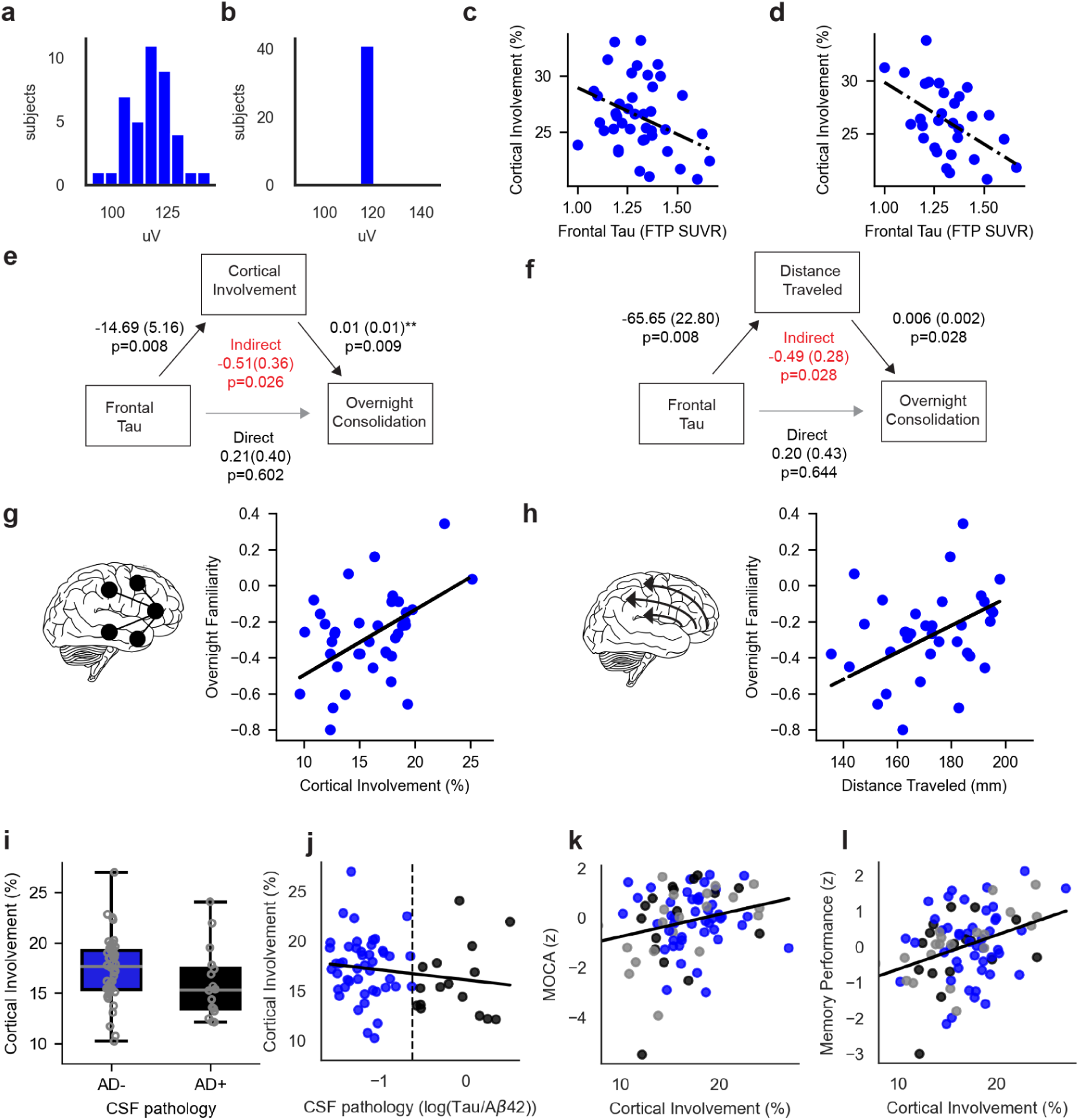
Slow Oscillation travel mediates the effects of frontal tau on memory consolidation. (a) Original distribution of slow wave amplitude across elderly participants (b) Distribution of amplitude across participants, after artificially removing slow waves **(c)** Scatter and resulting regression line of the amplitude-equated elderly population **(d)** Scatter and regression after sub-selecting the 1st 100 waves per participant to control differences in the number of waves per participant **(e)** Cortical involvement indirectly mediates the effect of frontal tau on overnight memory consolidation **(f)** Distance traveled by slow oscillations indirectly mediates the effect of frontal tau on overnight memory consolidation **(g)** Overnight familiarity is predicted by cortical involvement **(h)** Overnight familiarity is predicted by distance traveled by slow waves **(i)** Cortical involvement in AD+ participants is lower than cortical involvement in AD- in the BASE cohort **(j)** Cortical involvement is negatively correlated with AD pathology quantified by log(Tau/A*β*42) in the BASE cohort, AD- participants are colored in blue while AD- participants are colored in black, dashed line represents the level defined as AD positivity as in Fig 3i. (k) MoCA (Z-score) was positively correlated with cortical involvement in the BASE cohort, colors are the same while gray indicates biomarker data was unavailable **(l)** Memory performance (Craft Story, Z-score) is positively correlated with cortical involvement, colors are the same in 3k.

The same was true for the degree of impoverished cortical slow wave propagation. Controlling for chronological age, as well as sex, Apnea-Hypopnea Index (AHI), the degree of amyloid-beta burden (indexed using [^11^C]PIB), any time-difference between the tau and amyloid PET scans, as well as the degree of gray matter atrophy within the frontal ROI, the extent of frontal tau burden still significantly predicted the degree of impaired traveling propagation distance of slow wave across the scalp (β=-58.77 T=-2.59 p=0.02, r^2^=0.48; and see full table 2 in extended data, and **Fig 2h** for FDR corrected analysis of all areas).

**Table 2.** -Linear regression of frontal tau and distance travel.

| | $\beta$ | SE | T | p-value | CI[2.5%] | CI[97.5%] | Relative importance (%) |
| --- | --- | --- | --- | --- | --- | --- | --- |
| Intercept | 321.78 | 69.92 | 4.60 | 0.00 | 178.05 | 465.51 |  |
| Frontal Tau | -58.77 | 22.66 | -2.59 | 0.02 | -105.34 | -12.20 | 44.26 |
| Age | -0.10 | 0.50 | -0.20 | 0.84 | -1.12 | 0.92 | 2.76 |
| PiB | -4.44 | 15.88 | -0.28 | 0.78 | -37.08 | 28.19 | 10.50 |
| Sex (Male) | -9.74 | 5.53 | -1.76 | 0.09 | -21.11 | 1.63 | 11.14 |
| AHI | -0.55 | 0.28 | -1.96 | 0.06 | -1.13 | 0.03 | 27.62 |
| Frontal Thickness | -3.61 | 3.47 | -1.04 | 0.31 | -10.74 | 3.53 | 3.72 |

Having established that aging compromises these two dynamic features of NREM slow waves, and that tau in the frontal cortex is proportionally associated with these oscillation impairments, the third hypothesis prediction was tested, which aimed to determine whether tau-related impairments of these two contemporary slow-wave oscillations mediated, in part, the long-recognized association between tau burden and impaired episodic memory^4^.

As described in Methods, participants in the Berkeley cohort performed an episodic memory test, with encoding and an immediate test taking place before their in-lab polysomnography night of sleep recording, and then tested again the following morning^10,23^. The episodic recognition memory task quantified both familiarity and recollection memory (see Methods), with analyses focused on the former based on its established sensitivity to preclinical prodromal AD among the elderly^24,25^ and tau deposition specifically^26^.

Consistent with the hypothesis, the degree of impaired en masse coordinated slow-wave at night was associated with the degree of impaired overnight memory consolidation, such that the “lonelier” the slow waves, the greater the amount of overnight forgetting. This association remained significant when accounting for the cofactors of age, sex, amyloid-beta (global PET PIB), frontal atrophy, and AHI, again suggesting an age-independent, pathology-specific relationship (**Fig 3f**, β=0.04, T=2.70, p=0.01, r2=0.35, and see table 3 in extended data).

**Table 3.** Linear regression of cortical involvement and overnight memory consolidation.

| | $\beta$ | SE | T | p-value | CI[2.5%] | CI[97.5%] | Relative importance (%) |
| --- | --- | --- | --- | --- | --- | --- | --- |
| Intercept | -3.47 | 1.19 | -2.91 | 0.01 | -5.93 | -1.02 |  |
| Cortical Involvement | 0.04 | 0.01 | 2.70 | 0.01 | 0.01 | 0.06 | 52.98 |
| Frontal Tau | 0.21 | 0.40 | 0.53 | 0.60 | -0.62 | 1.05 | 3.75 |
| Age | 0.01 | 0.01 | 1.22 | 0.23 | -0.01 | 0.03 | 4.65 |
| PiB | -0.08 | 0.24 | -0.31 | 0.76 | -0.57 | 0.42 | 0.47 |
| Sex (Male) | 0.04 | 0.09 | 0.46 | 0.65 | -0.14 | 0.22 | 1.38 |
| AHI | 0.00 | 0.00 | -0.24 | 0.81 | -0.01 | 0.01 | 1.02 |
| Frontal Thickness | 0.11 | 0.05 | 2.14 | 0.04 | 0.00 | 0.22 | 35.74 |

Similarly, the extent of impaired cortical slow-wave traveling propagation predicted the degree of impaired overnight memory consolation, and as with the measure of lonely waves, these associations with impaired cortical traveling slow waves remained robust when accounting for the same cofactors of age, sex, amyloid-beta (global PET PIB), frontal atrophy, and AHI, again suggesting an age-independent, pathology-specific relationship (**Fig 3h**, β=0.01, T=2.21, p=0.04, r2=0.32 using the same covariates and see table 4 in extended data).

**Table 4.** Linear regression of distance traveled and overnight memory consolidation.

| | $\beta$ | se | T | p-value | CI[2.5%] | CI[97.5%] | Relative Importance (%) |
| --- | --- | --- | --- | --- | --- | --- | --- |
| Intercept | -4.61 | 1.59 | -2.90 | 0.01 | -7.91 | -1.32 |  |
| Distance Traveled | 0.01 | 0.00 | 2.21 | 0.04 | 0.00 | 0.01 | 36.88 |
| Frontal Tau | 0.21 | 0.44 | 0.47 | 0.64 | -0.70 | 1.11 | 3.59 |
| Age | 0.01 | 0.01 | 0.71 | 0.48 | -0.01 | 0.02 | 2.64 |
| PiB | -0.01 | 0.26 | -0.02 | 0.98 | -0.54 | 0.53 | 0.71 |
| Sex (Male) | 0.05 | 0.10 | 0.46 | 0.65 | -0.16 | 0.25 | 1.15 |
| AHI | 0.00 | 0.01 | 0.57 | 0.57 | -0.01 | 0.01 | 2.75 |
| Frontal Thickness | 0.15 | 0.06 | 2.62 | 0.02 | 0.03 | 0.27 | 52.28 |

Finally, the hypothesis tested was that tau-associated memory deficits in older adults are not simply direct, but partially mediated through tau-related impairments in the ensemble coordinated activity and traveling wave propagation. A bootstrapped mediation analysis revealed that the extent of diminished traveling wave propagation, and the degree of lonely, rather than ensemble coordinated, slow-wave activity mediated the effect of tau on overnight memory retention, predicting higher amounts of next-day forgetting, rather than remembering (cortical involvement indirect mediation: β = −0.051 (0.36), p = 0.02, 95% CI (−1.33 to −0.02); **Fig 3e**; and propagation distance traveled: β = −0.53, p = 0.02, 95% CI (−1.33 to −0.02); **Fig 3g**). Further, we tested whether the additional brain areas that predicted impaired cortical involvement (parahippocampal gyri and isthmus cingulate) as derived from whole brain FDR corrected regression analysis (**Fig 2h**) mediated the effect as well and found no significant mediation with the parahippocampal gyri (β = −0.21, p = 0.12, 95% CI [−0.65 to 0.05]) but a significant mediation with the the isthmus cingulate gyrus (β = −0.32, p = 0.04, 95% CI [−0.98 to 0.0006]) connecting to the parahippocampal gyrus in the medial temporal lobe, suggesting that the role of this brain area in slow wave travel and overnight memory consolidation should be further explored.

Next, slow waves were analyzed in an independent cohort, a subset of the BASE study at Washington University in St Louis, to address the replicability and generalizability of the results. Participants (Mean Age 70.82±4.57 years, U=5148 p=10^-6^) ranged in their cognitive status with an average MoCA score of 26.56±2.80 (range: 15-30) and a global CDR = 0.5 (very mild impairment) in 20% of the participants, and the remaining having a score of 0. In the morning, following overnight polysomnographic recordings, CSF amyloid and tau was derived using lumbar puncture. A total of 15 participants (18.5%) were classified as Alzheimer-Disease positive status (AD+) as determined by CSF [total tau/Aβ42] ratio.

EEG data from the BASE cohort was preprocessed and analyzed (see Methods) to assess both the degree of cortical involvement of the slow oscillations and the cortical traveling nature of slow waves as in the Berkeley cohort. On average, slow oscillations were co-detected in 16.85% ±3.62% of the EEG electrodes in the BASE cohort (similar to the Berkeley cohort T=1.73 p=0.08). With respect to travel, the average propagation across the scalp was 174.80mm±15.47 (and similar to the Berkeley cohort U=2887 p=0.35). The hypothesis posited that similar to the healthier Berkeley cohort, the degree of cortical involvement in slow waves and their propagation across the cortex will be diminished by chronological age. Beyond chronological aging, lower cortical involvement and diminished wave propagation will associated with higher levels of AD pathology in the CSF, and worse cognitive performance. The analysis demonstrated that higher age was correlated with lower cortical involvement (r=-0.37, [-0.55, -0.17], p=0.0005), and also diminished wave propagation (r=-0.41, CI=[-0.58, -0.2], p=0.002). In addition, despite the lack of spatial specificity of the lumbar puncture CSF measure, patients classified as AD+ (see methods) showed a trend for diminished cortical involvement of 16.0% ±.3.51% compared with AD- showing 17.4% ±.3.15% (U=484, p=0.03 confirmatory one-sided, **Fig 3i**), but wave propagation was not significantly different (170mm±16.50 in AD+ and 176mm±15.80 in AD-; U=417, p=0.08 confirmatory one-sided). In addition, higher levels of log[total Tau/Aβ42] were associated with low cortical involvement (N=62, Spearman r^2^=0.23, p=0.04 confirmatory one-sided, **Fig 3j**, while other CSF derivations were not significant, **Fig S4**).

Cognitively, participants with abnormal (<26) MoCA scores (N=22) showed significantly lower cortical involvement (15.2±3.91% compared to 17.5%±3.37% in those with normal MoCA score (≥26)]; U=404, p=.007) but with no difference in propagation distance (U=525, p=0.53). Similarly, MoCA scores were positively correlated with cortical involvement (r = 0.24, p = 0.02), however, the association was not significant at alpha = .05 after controlling for age, sex, and AHI (p = 0.055). Memory performance, which was assessed at a separate daytime visit before the polysomnographic recording, correlated with the degree of cortical involvement (N=77, r^2^=0.23, β=0.06, p=0.05, **Fig 3k**) after controlling for age and sex, and AHI (see table 5 in extended data).

**Table 5.** Linear regression of cortical involvement and memory performance in the BASE cohort.

| | $\beta$ | se | T | p-value | CI[2.5%] | CI[97.5%] | Relative Importance (%) |
| --- | --- | --- | --- | --- | --- | --- | --- |
| Intercept | -2.96 | 1.93 | -1.53 | 0.13 | -6.81 | 0.90 |  |
| Cortical Involvement | 0.06 | 0.03 | 2.03 | 0.05 | 0.001 | 0.13 | 33.86 |
| Age | 0.02 | 0.02 | 0.95 | 0.35 | -0.025 | 0.07 | 2.57 |
| Sex, Male | -0.61 | 0.20 | -3.03 | 0.003 | -0.21 | -1.02 | 59.46 |
| AHI | 0.01 | 0.01 | 0.82 | 0.41 | -0.01 | 0.02 | 4.10 |

**Table 6.** Linear regression of cortical involvement and MoCA in the BASE cohort.

| | $\beta$ | se | T | p-value | CI[2.5%] | CI[97.5%] | Relative Importance (%) |
| --- | --- | --- | --- | --- | --- | --- | --- |
| Intercept | -7.29 | 3.05 | -2.39 | 0.02 | -13.36 | -1.22 |  |
| Cortical Involvement | 0.10 | 0.05 | 1.95 | 0.05 | 0.00 | 0.19 | 32.10 |
| Age | 0.08 | 0.04 | 2.03 | 0.05 | 0.00 | 0.15 | 17.32 |
| Sex, Male | -0.65 | 0.32 | -2.03 | 0.05 | -0.01 | -1.28 | 41.83 |
| AHI | -0.02 | 0.01 | -1.28 | 0.20 | -0.04 | 0.01 | 8.75 |

Taken together, this collection of findings, across two cohorts and using two different measures of *in vivo* AD pathology, establish 1) that aging significantly impairs both the mass co-occurring, coordinated unity of slow waves throughout the brain, resulting in “lonely” slow waves, and further impairs the dynamic propagating, traveling nature of slow waves across the scalp, 2) the deficiency of these two novel properties of slow waves in older adults is age-related (i.e., evinced by the difference between young and older adults), but is not age-*dependent*, instead being most accurately explained by the pathological feature of tau protein accumulation in slow-wave generating regions of the brain, 3) the degree of impairment in both the en masse unity of slow-wave occurrence and the diminution in traveling cortical propagation predict the extent of failed overnight memory consolidation in the elderly, associated with a failure of overnight memory retention, and 4) the tau-associated overnight memory consolidation deficits are not direct, but instead, partially mediated through impairments in these two sleep oscillation features of ensemble coordinated activity and traveling wave propagation.

Previous studies have demonstrated that aging is accompanied by a decline in the synchronization of brain activity across cortical regions, and is linked to diminished executive and memory performance^27^. The current study establishes that slow oscillations in NREM sleep, which are otherwise the most synchronized and unified events in the healthy brain, can become disunited and thus isolated in the aging brain. Notably, this phenomenon was not solely attributable to a reduction in cortical thickness associated with aging, but rather to the accumulation of tau protein, particularly evident in frontal gyri and select regions within the medial temporal lobe. Areas whose communication during sleep was associated with overnight memory consolidation^28^. Additionally, high levels of AD pathology in the CSF, while lacking spatial specificity, were also predictive of isolated slow waves. However, this effect was not as robust and specific to cortical involvement suggesting that the accumulation of tau, specifically in the frontal and some medial temporal regions, is disruptive to achieving slow wave function in overnight memory consolidation.

How tau disrupts neural activity and slow oscillations is not clear. Recent studies show that it is not the NFTs imaged by FTP-PET but the associated soluble tau stemming from the breakage of microtubules that suppress neural activity^29,30^. Injection of soluble tau into single pathology-free neurons in a slice takes 30 minutes to blunt action potentials, decrease their amplitude, and slow their rise, resulting in reduced excitatory post-synaptic potential (EPSP) in connected neurons^31^. Studies investigating the effect of tau on neural networks in-vivo are conflicted. While the evidence in support of hyperexcitability due to the accumulation of amyloid is strong, it is not clear if on the network level tau is also causing hyperexcitability or vice versa^32^. The current study shows that the accumulation of tau in the medial-temporal-lobe-frontal network of humans is associated with decreased excitability, as slow oscillations do not propagate, resulting in lonely slow waves. Therefore, while tau accumulation might be a result from a hyperexcitable neural network^33^, it seems that its accumulation acts in the opposite depressive direction, at least during sleep, and similar to observations in sleeping and anesthetized mice^34,35^. Since impaired slow waves are observed early, in unimpaired elderly with sub-clinical memory impairment, it could constitute a complimentary early detection method to assess neurodegeneration levels, especially if further confirmed longitudinally and in patients.

What is the function of global traveling slow waves? Not all slow waves are collective or traveling, as they can occur locally after region-specific learning^36^, during wakefulness after sleep deprivation^37,38^, or restricted to a single hemisphere^39^, yet many slow waves do encompass large cortical territories. Do these system-level events differ in their function? This study shows, for the first time, that losing these wide-reaching propagating slow waves is detrimental to overnight memory consolidation, suggesting that coordinated quiescence across the cortex plays a role in adequately modifying brain-wide neural networks overnight. A sequential calcium influx following electric slow oscillations could be modifying neural populations serially^13^. In wakefulness, sequential activation of a neural network enables sequential information processing, while later active populations have access to already generated outcomes^40^. During information processing frontal areas tend to respond last, hundreds of milliseconds after sensory areas in a feedforward sweep^41^. A reverse flow is evident in traveling slow oscillations, similar to back-propagation of the error signal required for updating in artificial neural networks. Coordinated propagating slow oscillations could underlie an ability to update a brain-wide network offline, according to the events of the day into a declarative schema.

## Methods

### Participants

The Berkeley cohort included a total of 138 participants. Out of them, 61 were young healthy adults (mean age=20.24±1.91 years, 49% males). Older adults were cognitively normal individuals recruited from the Berkeley Aging Cohort Study (BACS) (N=74, mean age= 74.79, SD= 5.73, 32% male). Data from most of these participants was included in previous publications^10,23,42–45^. Exclusion criteria were a history of neurologic, psychiatric, or sleep disorders, current use of antidepressant or hypnotic medications, and Mini-Mental State Examination score (MMSE) <25^46^.

The Biomarkers of Alzheimer Disease in Sleep and EEG (BASE) cohort reported here (N=82, mean age: 70.5±4.4 years, 48% male) is a subset of the BASE study at Washington University in St. Louis. The BASE study enrolls community-dwelling participants aged ≥65 years from the St. Louis, MO region. Exclusion criteria were known neurological disorder or brain injury, CDR of ≥ 2, BMI > 40 kg/m2, and contraindication to lumbar puncture.

### General Experimental Procedure

Older Participants in the Berkley cohort underwent structural 1.5T MRI, tau PET with 18F-Flortaucipir (FTP), Aβ PET with 11C-Pittsburgh compound-B (PiB). Within 1.5 years of PET scanning, participants completed two sleep study sessions separated by a week or two. On both nights, participants were given 8-hour sleep opportunities monitored with polysomnography (PSG, see below). The second night served as the experimental night, and on that night the participants were presented with the paired association learning task (see details below). In the morning after, high-resolution structural MRI scans were obtained (see details below) from all participants to measure gray matter atrophy. Subsequently, participants completed the retrieval blocks of the memory task. All participants abstained from caffeine, alcohol, and daytime naps for 48 hours before and during the sleep study sessions. Participants kept habitual sleep-wake rhythms for at least 1 week preceding the in-laboratory sleep session and completed the in-lab sleep study in accordance with their habitual bedtime. Young participants in the Berkeley cohorts underwent different experimental procedures as described in previous publications (N=19^23^, N=25^47^, and N=17^48^).

Participants in the BASE cohort at Washington University **c**ompleted an initial onsite visit consisting of clinical evaluation and psychometric testing, including the Montreal Cognitive Assessment^49^ (MoCA, general cognition), Craft Story 21 recall immediate and delayed (memory). Clinical Dementia Rating (CDR) scores were determined by clinical evaluation^50^. Within 5-14 days of their initial visit, participants completed an in-lab overnight full polysomnography (see below). The following morning (9-10AM), cerebrospinal fluid (CSF) was obtained by lumbar puncture (see below).

### Sleep monitoring

Polysomnography of the Berkeley cohort was recorded using a 19-channel EEG Grass Technologies Comet XL system (Astro-Med, Inc., West Warwick, RI, USA) and digitized at 400Hz. Polysomnography of the BASE cohort was recorded using a 20-channel EEG via a hard-wired, dedicated Grael 4K PSG:EEG (Compumedics, Victoria, Australia) system and Profusion Sleep 4™ software and digitized at 256Hz.

In both cohorts, EEG electrodes were placed according to the 10–20 system and referenced to contralateral mastoids. Electrooculography (EOG) was recorded at the left inferior and right superior outer canthi, and electromyography (EMG) on the chin.

### EEG analysis

In both cohorts, sleep scoring was performed according to the guidelines of the American Association of Sleep Medicine^51^ and blinded to pathology levels. Then, EEG was preprocessed using MNE-python package^52^. Visually obvious artifact-free EEG Data scored as either N2,N3 sleep was extracted from all subjects, bandpass filtered between 0.5-40Hz using a finite impulse response (FIR) filter, and downsampled to 200Hz. Then, data and further subjected to a threshold of 400μv to exclude artifacts on a 5sec window basis. Overall, this procedure resulted in an average of 93±6% of the data for young participants available for further analysis and an average of 97±10% in the older participants in the Berkeley cohort and 99±1% in the older adults in the Washington cohort. Lower oscillatory power in the slow wave (<4Hz) range was observed for older adults as previously reported^42^ (**Fig S1**), and therefore analysis futher focused on the detection of individual slow oscillations.

Individual slow wave oscillation events were detected (slow oscillations, 0.3–1.5 Hz) using established algorithms^11,18^ as implemented in the YASA python package^17^. In short, each electrode was bandpass between 0.3-1.5 using a FIR filter with a transition band of 0.2Hz. Negative peaks between -40 and -200μv were detected and positive peaks between 10 and 150μv. For each negative peak (slow wave trough), the nearest positive peak was found and the peak-to-peak amplitude was computed. In addition, the duration of the negative phase and positive phase was computed. The negative phase is the time between the first zero-crossing (before the negative peak) and the second zero-crossing (between the peaks) and the positive peak is the time between the zero-crossing between the peaks and the last zero-crossing (after the positive peak). Waves were included only if peak-to-peak amplitude was within 75-300μv, the negative phase was between 0.1-1.5sec and the positive peak was between 0.1-1sec.

Next, an analysis of cortical involvement and distance traveled was conducted, inspired by previous publications^53,54^ yet simplified and adapted for a clinical, low-density EEG. Slow wave detections were sorted according to the negative peak in 10ms resolution, and within that range according to the amplitude of the negative peak. The earliest channel detecting a valid negative peak was considered an origin, and additional electrodes detecting negative peaks before the mid-crossing of the previous wave were considered part of the same multichannel slow oscillation.

A negative peak detected after a previous wave mid-crossing was considered a new slow oscillation. After this grouping, 2 older subjects that showed less than 25 multichannel waves were excluded from the Berkeley cohort (resulting in N=74). Then, the degree of cortical involvement for a particular slow oscillation was the number of detecting electrodes divided by the number of EEG electrodes in the 10-20 montage (19 in both cohorts) to allow comparison across different systems and montages.

Both in older and young adults slow waves mostly originated in the frontal channels fitting prior high-density EEG studies^11^, EEG source analysis^15^, and intracranial recordings^14^. Five frontal channels (26%) were the origin of 56.61±7.88% of the waves and 68.72±13.34% of the waves in the older (U=3721,p<10^-8^). Propagation analysis was performed for waves with more than 2 participating electrodes and reapplying the threshold above for a minimal number of waves (Excluding 7 participants from the Berkeley cohort and 4 participants from the BASE Cohort from this analysis). Distances were calculated as the maximal distance between the electrodes detecting the same slow oscillations^53^ and using the standard locations in the MNE package.

Stratification analysis was carried out on each participant’s waves, by artificially excluding waves from the side of the amplitude distribution such that the median amplitude of the participant is as the median amplitude of the group (120µV) such that the entire population of older participants with various slow amplitude (**Fig 3a**) will have the same median amplitude (See **Fig 3a**) and recalculating cortical involvement and the correlation it with frontal tau (See **Fig 3c**). To control for differences in the number of waves, the correlation with tau was recalculated using only the first 100 waves per participant (**Fig 3d**).

### PET acquisition and processing

PET data acquisition was described previously^55–57^. Both FTP and PiB were synthesized at the Biomedical Isotope Facility at LBNL and all PET imaging was conducted on a BIOGRAPH PET/CT scanner, with FTP scans usually conducted on the same day immediately following PIB. For FTP scans, participants were first injected with 10 mCi of tracer, and data acquired from 80 to 100 min postinjection were used for analysis. CT scans collected before the start of emission acquisition were used for attenuation correction. We reconstructed the FTP-PET images using an ordered subset expectation maximization algorithm with scatter correction and smoothed with a 4-mm Gaussian kernel. For FTP data processing, the mean tracer retention over 80–100 min postinjection was normalized by the mean tracer retention in the inferior cerebellar gray, as the reference region, to create FTP standardized uptake value ratio (SUVR) images. We performed partial volume correction (PVC) to account for partial volume effects related to atrophy and spillover signal, using the Rousset geometric transfer matrix method, as detailed previously^58,59^. The frontal region of interest was defined according to Desikan Killany^60^ and included the frontal lobe areas including the superior frontal gyrus and both the rostral and casual division of the middle frontal gyros as well as the rostral and caudal cingulate cortex (see **Fig S5**).

For PiB-PET imaging, participants were injected with 15 mCi of PiB tracer, and 90 min of dynamic acquisition frames began immediately after the injection. A CT scan was obtained before the injection and used for attenuation correction. PiB-PET images were also reconstructed using an ordered subset expectation maximization algorithm with scatter correction and smoothed with a 4-mm Gaussian kernel. For PiB data processing, distribution volume ratio (DVR) was generated with Logan graphical analysis^61,62^ on frames over 35–90 min post-injection and normalized using the whole cerebellar gray as the reference region. Global PiB was calculated using multiple FreeSurfer ROIs across the cortex, as previously described^63^.

### Lumbar Puncture

Fasted cerebrospinal fluid (CSF) was collected at 9-10 AM the morning following PSG by lumbar puncture and immediately placed on ice, prior to centrifugation and storage at -80C. Amyloid-β40 (Aβ40), amyloid-β42 (Aβ42), total tau, and phosphorylated tau 181 were assessed by Lumipulse by the Washington University Knight Alzheimer Disease Research Center Biomarker Core. The ratio of total tau to Aβ42 was used to determine AD status, with those having [total tau/AβB42] > 0.541 categorized as AD+^64^. Currently, the biomarker results from N=62 were available for analysis.

### Memory Assessment

**Participants in the Berkeley cohort** were assessed using an overnight memory task. In the evening, before the overnight sleep recording, older participants learned the association between pairs of words and non-words (paired association task) until a criterion was met (see^10,23^ for details). Then, in the morning, participants were tested using a forced choice (new, learned-pair, lure) and analyzed to create two scores: familiarity accuracy and recollection. Paired associative learning (PAL) had been previously proven sensitive to the effect of sleep^65–69^ and especially familiarity was previously shown to be sensitive to pathological aging^24,25^.

Familiarity score was defined as:

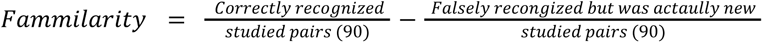

As opposed to recollection:

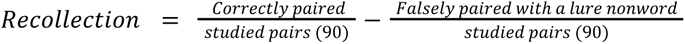

Overnight familiarity memory change was quantified by subtracting the evening accuracy from the morning accuracy.

**Participants in the Washington cohort** were assessed using Craft Story 21 recall immediate and delayed (memory)^70^ collected during an afternoon visit 1-2 weeks before the overnight PSG. In brief, the Craft Story 21 Recall evaluates the capacity to remember a short narrative. The examiner articulates the story to the participant in a clear manner. Following the narration, the participant is tasked with recollecting the story from memory without any repetitions allowed. Subsequently, after a 20-minute delay, the participant is prompted to reproduce the story, with cues provided to enhance recollection for later assessment. Memory performance was z-scored, and adjusted for age, sex, and education using the multi-domain impairment score (MIRS)^71^ as implemented in the online Mayo Clinic calculator^72^. The Craft Story 21 recall immediate (VRS) and delayed (DVR) variables were highly correlated in this cohort (r=0.83) and were thus averaged prior to analyses.

### MRI scanning and analysis

Structural MRI scans were acquired in order to assess brain atrophy. All participants in the Berkeley cohort were scanned in a Siemens Trio 3T scanner right after the sleep session. High-resolution anatomic images were collected using a T1-weighted magnetization-prepared rapid gradient-echo (MPRAGE, Repetition Time (TR) = 1900 echo time (TE) =2.52 ms, Flip Angle (FA) = 9°, 1mm isotropic voxels). The scans were processed using FreeSurfer version 5.3^60,73,74^. Briefly, after co-registering each image to MNI305^75^, FreeSurfer reconstructs three-dimensional (3D) pial and white matter surfaces, based on the relative intensity differences at the boundaries of white and gray matter tissue. Cortical thickness (in mm) was calculated across≈150,000 vertices per hemisphere as the average distance of the vectors perpendicular to the triangular faces of the white matter and pial surfaces^76^. Then, cortical parcellation was performed based on regions of interest (ROIs) as defined by the Desikan-Killiany atlas^60,73^. The frontal region of interest used for the regression was derived by averaging the Desikan Killany Atlas frontal areas described above for the tau-PET data.

### Statistical analysis

All statistical comparisons were carried out using the Pingouin Python package for statistical analysis^66^. All correlations reported are Pearson correlations unless stated otherwise. Linear regression was performed using the same Pingouin package as well as mediation analysis using bootstrapping (All using 1000 iterations). Direct statistical comparison analysis was carried out using a t-test when both populations were normally distributed (verified with Shapiro-Wilk test) and using a Mann-Whitney test otherwise, and then a U was reported instead of a T value and the criteria for significance was 0.05 two-sided unless stated otherwise. Multiple-comparisons correction was performed using Benjamini-Hochberg method for false discovery rate (FDR)^77^ as implemented in Python stats models package^78^.

## Ethics approval and consent to participate

Berkeley Cohort

The study was conducted in accordance with the Declaration of Helsinki and approved by the human studies committees at the University of California, Berkeley and Lawrence Berkeley National Laboratory (Berkeley Committee for Protection of Human Subjects Protocol Numbers 2010-01-595 and 2015-03-7268), with all participants providing written informed consent.

BASE Cohort

The Biomarkers of Alzheimer’s Disease in Sleep and Electroencephalography (BASE) study was approved by the Human Research Protection Office at Washington University in St. Louis (Protocol 201804006), and all participants provided written, informed consent.

## Data availability

All source data for the main figures and Extended Data figures will be made available upon publication.

## Code availability

Python scripts to calculate cortical involvement and slow wave propagation will be made available upon publication.

## Supplementary materials

**Fig S1.**
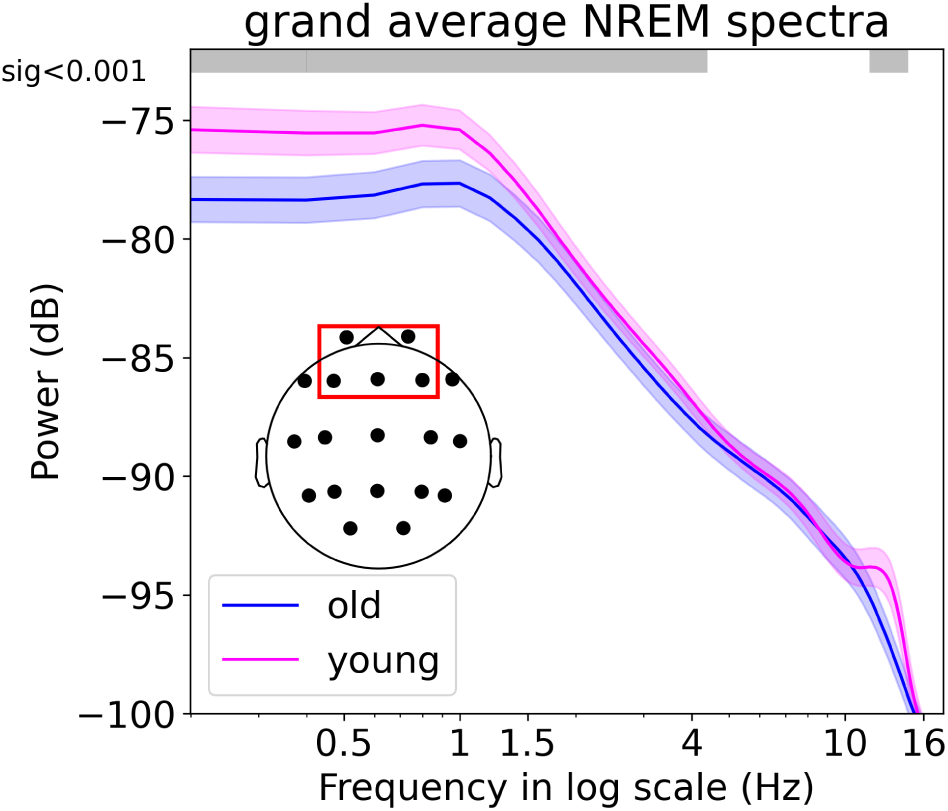
the spectral difference between young and older adults in frontal channels. Spectral power using multi-taper Fourier analysis on average 5 frontal electrodes (see inset) in young (magenta) and older adults (blue) presented in logarithmic scale. The gray bar above indicates the significance value of p<0.001 after FDR correction (see methods).

**Fig S2.**
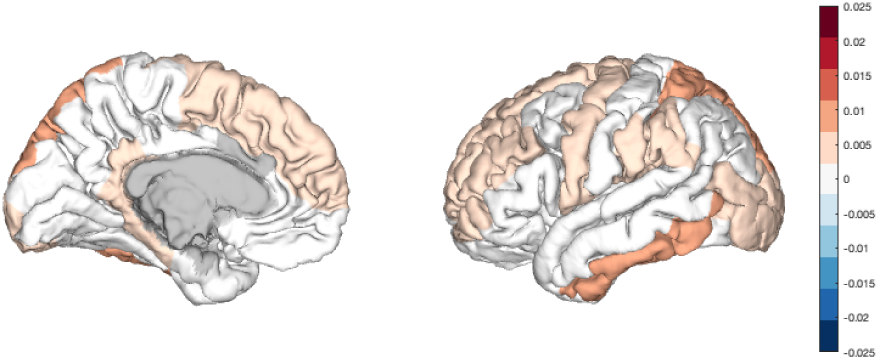
Apnea-Hypopnea Index (AHI) and accumulation of tau. Apnea-Hypopnea Index (AHI) had significant but mildly predictive power on tau levels in frontal ROI beyond age and sex (β= 0.005, p=0.02, r=0.13). Nevertheless, this effect was not spatially specific and apparent in various brain regions and was included as a covariate in the main analysis above.

**Fig S3.**
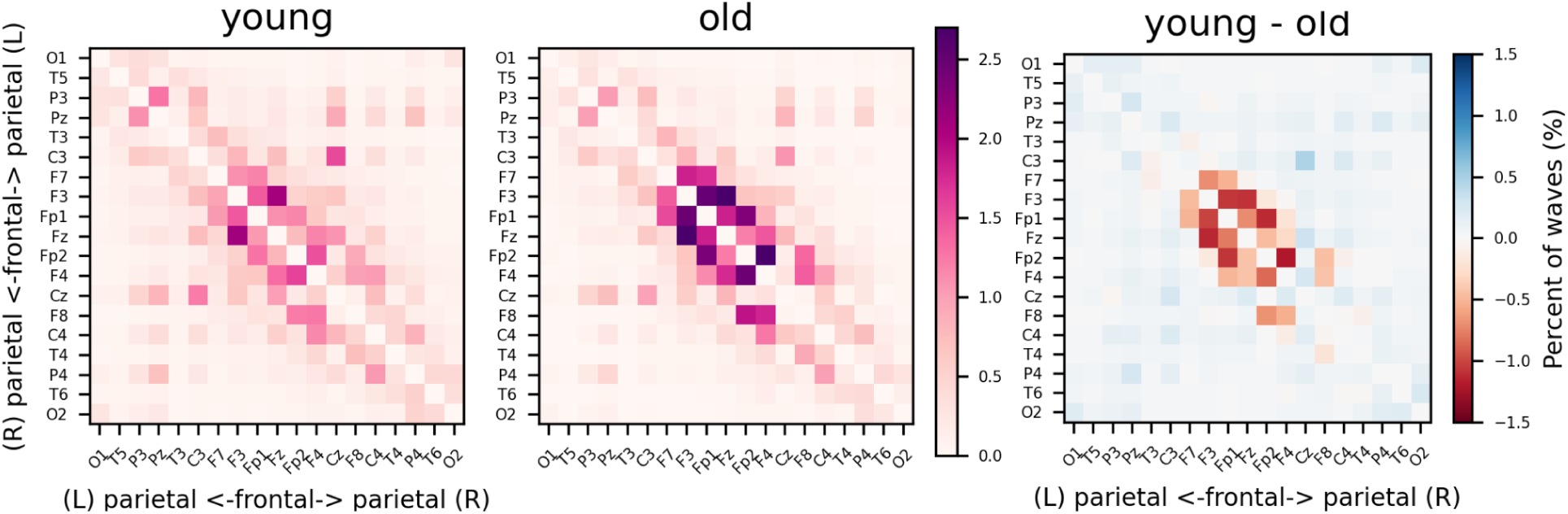
Slow travel in relative terms. Left, the average percent of slow oscillations traveling between each pair of the electrodes in the young participants, the center of the matrix represents frontal electrodes while the upper part represents the left hemisphere from the front to back and the lower part the right hemisphere in the same manner (see Figure 2b). Center, the same figure for old participants. Note how in relative terms, the slow oscillations of the older participants tend to be concentrated in frontal areas while in young the there is more relative travel detected in partial electrodes. Right, the difference between young and older adults groups.

**Fig S4.**
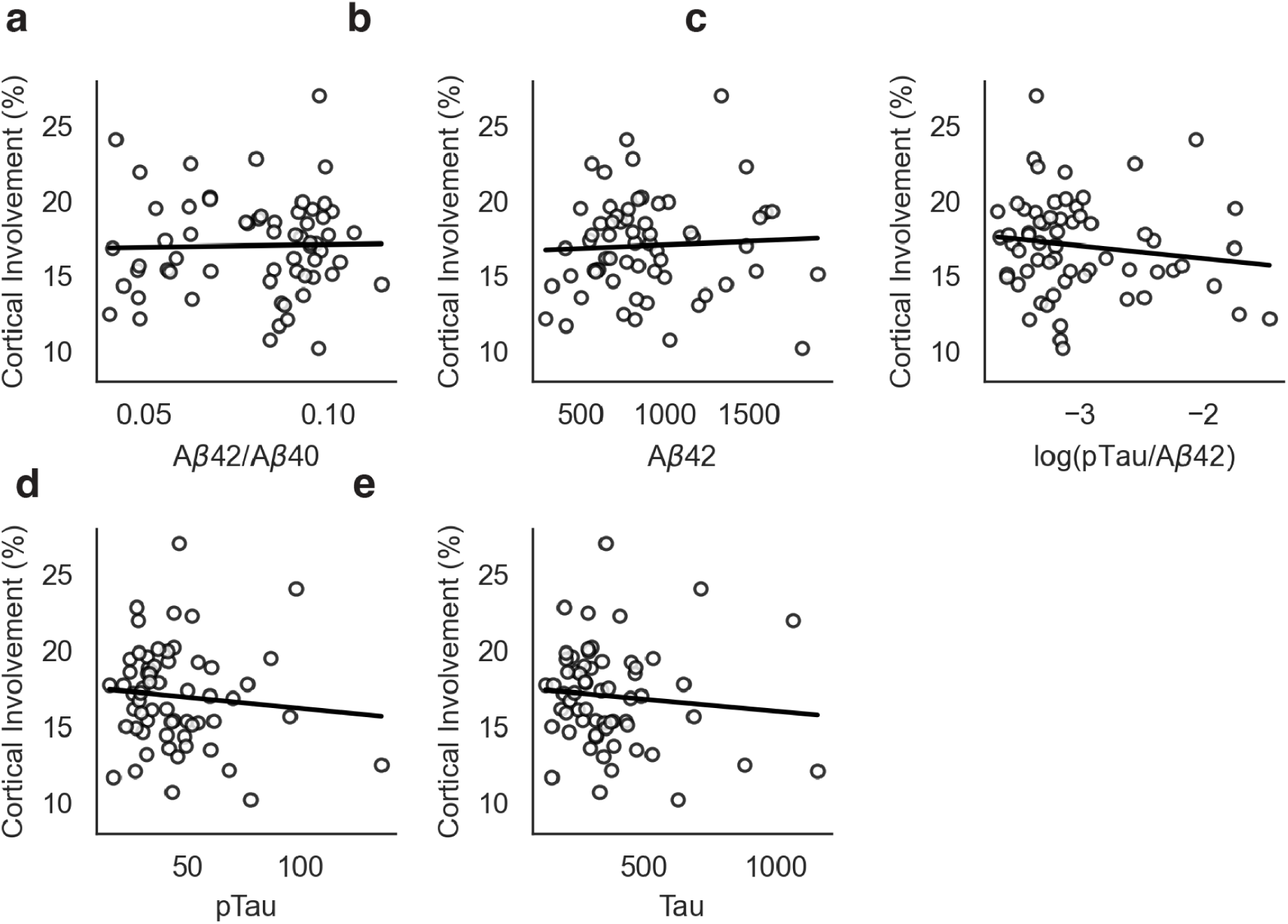
Cortical involvement and additional CSF derivatives. Cortical involvement and CSF biomarkers. a) Ratio of amyloid-beta42 to amyloid-beta40 b) Amyloid-beta42 c) Ratio of phosphorylated tau to amyloid-beta42, log-transformed d) Phosphorylated tau e) Tau. None of the associations were significant (p > .05 for all).

**Fig S5.**
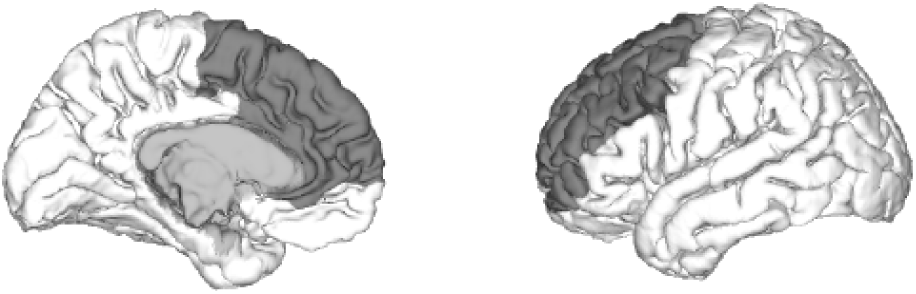
the predefined frontal region of interest. Areas used as a-priori region of interest, defined according to Desikan Killany atlas and included the frontal lobe areas including the superior frontal gyrus and both the rostral and casual division of the middle frontal gyros as well as the rostral and caudal cingulate cortex

## Acknowledgments

This work was supported by NIH grant RF1AG054106 awarded to M.P.W.

## Notes

### Competing Interest Statement

The authors have declared no competing interest.

